# Prediction of Interactions between Cell Surface Proteins by Machine Learning

**DOI:** 10.1101/2023.09.12.557337

**Authors:** Zhaoqian Su, Brian Griffin, Scott Emmons, Yinghao Wu

## Abstract

Cells detect changes of external environments or communicate with each other through proteins on their surfaces. These cell surface proteins form a complicated network of interactions in order to fulfill their functions. The interactions between cell surface proteins are highly dynamic and thus challenging to detect using traditional experimental techniques. Here we tackle this challenge by a computational framework. The primary focus of the framework is to develop new tools to identify interactions between domains in immunoglobulin (Ig) fold, which is the most abundant domain family in cell surface proteins. These interactions could be formed between ligands and receptors from different cells, or between proteins on the same cell surface. In practice, we collected all structural data of Ig domain interactions and transformed them into an interface fragment pair library. A high dimensional profile can be then constructed from the library for a given pair of query protein sequences. Multiple machine learning models were used to read this profile, so that the probability of interaction between the query proteins can be predicted. We tested our models to an experimentally derived dataset which contains 564 cell surface proteins in human. The cross-validation results show that we can achieve higher than 70% accuracy in identifying the PPIs within this dataset. We then applied this method to a group of 46 cell surface proteins in C elegans. We screened every possible interaction between these proteins. Many interactions recognized by our machine learning classifiers have been experimentally confirmed in the literatures. In conclusion, our computational platform serves a useful tool to help identifying potential new interactions between cell surface proteins in addition to current state-of-the-art experimental techniques. The tool is freely accessible for use by the scientific community. Moreover, the general framework of the machine learning classification can also be extended to study interactions of proteins in other domain superfamilies.

## 1. Introduction

The plasma membrane provides the structural support for cells and protects them from the outside environment [1]. Along the surface of the plasma membrane, different species of proteins are arranged, collectively referred to as cell surface proteins. Cells communicate with each other and receive external signals through these molecules [2, 3]. Many cell surface proteins are thus important targets for biomedical research due to their accessibility for pharmacological intervention and utility as biomarkers [4]. Most of them contain an extracellular region, one or multiple trans-membrane segments which are embedded in the plasma membrane, followed by a cytoplasmic region [5]. Spanning across the plasma membrane, cell surface proteins carry out their cellular functions by interacting with each other [6–9]. For instance, during cell communications, the extracellular region of a cell surface protein interacts with the extracellular regions of others on neighboring cells [10, 11]. The messages of these extracellular interactions are transferred to the cytoplasmic regions of the corresponding proteins so that the intracellular signaling pathways will be activated [12–14]. One typical example is the formation, maturation, and plasticity of synapses between neurons, which are closely maintained by the spatial-temporal interactions among different cell surface proteins such as Down syndrome cell adhesion molecules (DSCAMs) [15]. In another case, immune responses are tightly regulated by the interactions between a large variety of cell surface receptors such as programed cell death-1 (PD-1) and its ligand programed cell death-ligand 1 (PD-L1) [16]. As a result, a thorough study on a complete list of interactions between all cell surface proteins can significantly facilitate our understanding to their functions in regulating cell communications and signaling.

It is currently challenging to measure the extracellular interactions between cell surface proteins on a systematic level [17]. These interactions are difficult to detect by standard biochemical assays due to the transient nature of their binding kinetics which spans a broad range of affinities from low nM to hundreds of mM [18]. Although genome-wide networks of protein-protein interactions (PPIs) have been constructed for many species based on the technological improvements in high-throughput experiments such as affinity-mass spectrometry [19] and yeast-two-hybrid [20], these approaches have been proven effective in identify binding between cytoplasmic proteins but inadequate to discover interactions among proteins that are associated with the plasma membrane [21]. On the other hand, computational modeling not only serves as an ideal alternative approach to complement time-consuming and labor-intensive experiments, but also can test conditions that are currently inaccessible in the laboratory. A large variety of methods have been developed to predict PPIs [22, 23]. Predictions in these methods rely on different types of data source, include phylogenetic profiles [24], literature mining [25], gene ontology annotations [26], co-evolution of interacting proteins [27], etc. Additionally, the new advancements in machine learning and deep learning have had huge impacts on the field of computational biology [28, 29]. The application of these algorithms to PPIs has gained enormous successes [30, 31]. However, the models in these studies were trained by more generalized datasets of PPIs. They are not sensitive enough to recognize the specific interactions between cell surface proteins. As we will show in the results below, interactome of cell surface proteins predicted from these methods has little overlap with the experimentally derived one. A machine-learning-based prediction method that focuses on the extracellular interactions between cell surface proteins, to the best of our knowledge, is still lacking.

In this study, we designed a machine-learning-based computational framework to predict interactions between cell surface proteins. Our method specifically focuses on proteins with domains in the immunoglobulin (Ig) fold, which are called Ig domains hereafter. Ig domains share the structural characteristics of a β-sandwich framework [32], and can form homotypic or heterotypic interactions with each other [33]. They are the most abundant domain family functioned for cell surface recognition, and are widely distributed in a large portion of cell surface proteins involving in cell adhesion and signaling [34]. For example, the extracellular region of DSCAM contains 16 Ig domains. Some of these domains can form homophilic binding, thus regulating the interactions between DSCAMs from different neurons [35]. Similarly, the extracellular regions of most cell surface receptors and ligands involved in immune signaling, including PD-1 and PD-L1, also contain multiple copies of Ig domains. Their interactions lead to the formation of co-regulatory complexes at the interface between T cells and antigen-presenting cells (APC), which play a pivotal role in activating immune responses [36].

In practice, we constructed a non-redundant structural database of interactions between Ig domains. We collected all pairs of fragments located at the binding interfaces between Ig domains in the database. Based on this interface fragment pair library, we can predict if two cell surface proteins can form interaction or not. Given a pair of Ig domain proteins which interaction is unknown, we align their sequences to all the interface fragment pairs. The list of top alignment scores was used as indicators of multiple machine learning algorithms to calculate the probability of interaction. We tested our models to an experimentally derived dataset which contains 426 interactions among a total number of 564 cell surface proteins in human. The cross-validation results show that we can achieve higher than 70% accuracy in identifying the PPIs within this dataset. After calibrated our method against the human dataset, we further applied it to systematically predict the interactions of cell surface proteins in other species, namely, the nematode Caenorhabditis elegans (C. elegans). The binding properties of cell surface proteins in C. elegans is of great interest in the context of synaptic specificity. In C. elegans, all the synaptic connections in their nervous system are known [37]. However, it is unclear how these connections are genetically encoded. The cell surface proteins in C. elegans are thought to be important labels for cell-cell recognition in synapse formation. The cell-specific expression of these proteins in the nervous system is known from several studies [38]. If it could be further determined which pairs of these proteins interact, this would represent an important step in understanding the code for neuronal wiring. To reach this goal, a group of cell surface proteins from C. elegans which contain Ig domains in their extracellular regions were selected. We screened every possible interaction between these proteins. Most of the interactions recognized by our machine learning classifiers among a small subset of five well-studied proteins can be validated by previous experimental measurements in the literature. In summary, our computational method provides a useful tool to study interactions of cell surface proteins, while the results from our prediction can further help us understanding their functions.

## 2. Results

### 2.1. Construct the interface fragment pair library from a non-redundant structural database of interactions between Ig domains

We first constructed a non-redundant structural database of interactions between domains that belong to the Ig fold. Most domains in Ig-fold belong to two Pfam clans. The first is the immunoglobulin superfamily (CL0011), while the other one is the superfamily of immunoglobulin-like fold (CL0159). A detailed definition of domain-domain interactions and the procedure of database construction is described in the **Materials and Methods**. As a result, the database contains 831 pairs of domain-domain interactions. The structure of each interaction was then obtained from the Protein Databank (PDB). Some specific examples in our database are plotted in **Figure 1a**, with PDB IDs marked beneath their corresponding structures. Two interacting domains in the plot are colored in red and green, respectively. For instance, 1EBA is from an extracellular dimer of erythropoietin receptor, a member in the cytokine receptor family, responsible for preventing erythroid progenitors from apoptosis [39]. In another case, 3PPE shows the interactions between two N-terminal domains of VE-cadherin, which is a major endothelial adhesion molecule regulating blood vessel formation [40].

**Figure 1:**
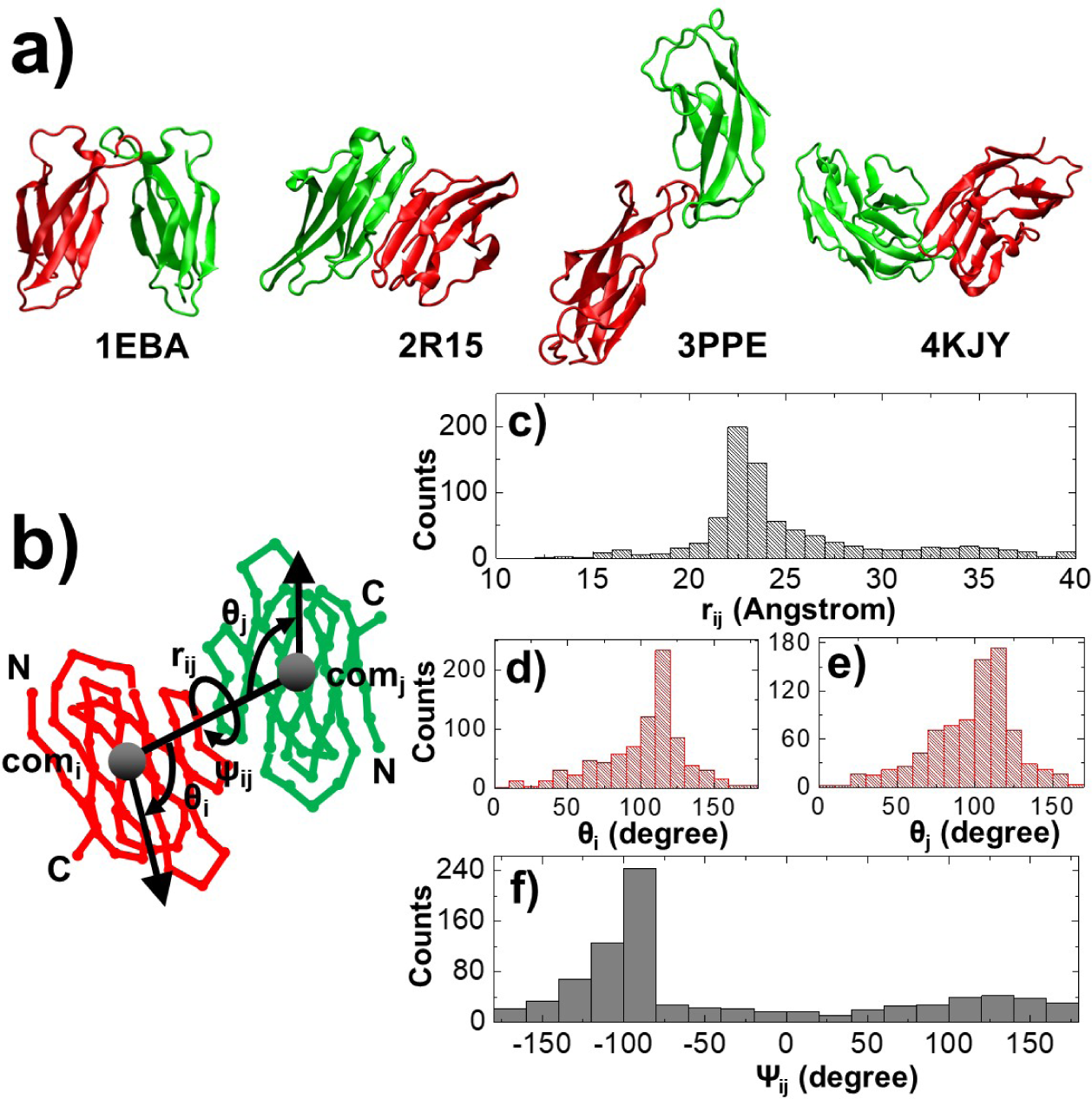
The structures of some Ig domain interactions from our non-redundant database are shown in **(a)**. Two interacting domains are colored in red and green, respectively. The corresponding PDB id is listed below each interaction. Given a specific pair of interacting domains, the conformation of their relative orientation is defined by four parameters: *r_ij_*, *θ_i_*, *θ_j_*, and *Ψ_ij_*, as shown in **(b)**. We calculated the values of these conformational parameters for all 831 domain pairs in the database. In detail, the distribution of distance between two domain centers of mass *r_ij_* is plotted as histograms in **(c)**. The distributions of two packing angles between domains *θ_i_* and *θ_j_* are plotted in **(d)** and **(e)**, respectively. Finally, the distribution of the tortional angle *Ψ_ij_* is plotted in **(f)**.

The global orientation of inter-domain binding between two Ig domains can be characterized by a minimal number of four conformational parameter, as described in **Figure 1b**. The first conformational parameter, *r_ij_*, is the distance between the centers of mass of two interacting domains. The next two parameters, *θ^i^* and *θ_j_*, define the packing angles between *r_ij_* and the vectors which point from the N-terminal to the C-terminal of each domain, respectively. The last parameter *Ψ_ij_* indicates the tortional angle between two domains. In order to explore the structural properties of binding between Ig domains, we perform statistical analysis to all interacting domains in the database. Specifically, for each of the 831 domain pairs, we calculated their conformational parameters based on above definition. In order to test whether the binding between Ig domains has any conformational preference, we plotted the distribution of each calculated parameter. **Figure 1c** shows the distribution of distance between two domain centers of mass. The figure indicates that the distance between the centers of mass of two interacting Ig domains is mostly within 20 to 25 Angstrom. Following the packing distance, **Figure 1d** and **1e** further show the distributions of two packing angles. The similarity of distributions in these two plots suggests the symmetry of binding between two domains. Moreover, the figures indicate that both interacting domains favor the packing angle around 120°. Finally, the distribution of the tortional angle is plotted in **Figure 1f**. The angle spans a wide range from -180° to 180°, although a peck at -90° was observed. Overall, the anisotropic distributions of different conformational parameters imply that the binding between Ig domains is not stochastically formed, but has certain level of preference.

In our previous study, we showed that the space of binding interfaces between proteins can be decomposed into smaller fragments given the fact that structural space of PPIs is highly degenerate [41]. We therefore hypothesized that this fragment-based representation of protein-protein binding interfaces has the potential in predicting and modeling new PPIs. Following this hypothesis, we highlighted the fragments located at binding interfaces of all interacting Ig domains in our constructed database, which are called interface fragment pairs. More specifically, an interface fragment pair is defined as a pair of peptide fragments with the length of 5 residues (**Figure 2a**). The first fragment is from one of the two interacting Ig domain, while the second fragment is located in the other interacting Ig domain. Previous studies suggested that a 5-residue fragment is an ideal length to appropriately capture the local secondary structure types, while being short enough to model the space of protein-protein binding interfaces with a tractable number of representative conformations [42]. Moreover, the sidechain of the two residues in the middle of both fragments should form atomic contacts with each other, as illustrated by the enlarged insertion of the figure. The distance cutoff used is assigned to 5 Angstrom. This distance cutoff was adopted from previous studies [43] to define the contacts between heavy atoms of the sidechains from two interacting proteins. Given these criteria, we scanned the structures of all 831 non-redundant Ig domains interaction. As a result, a total number of 37968 interface fragment pairs were collected from our database.

**Figure 2:**
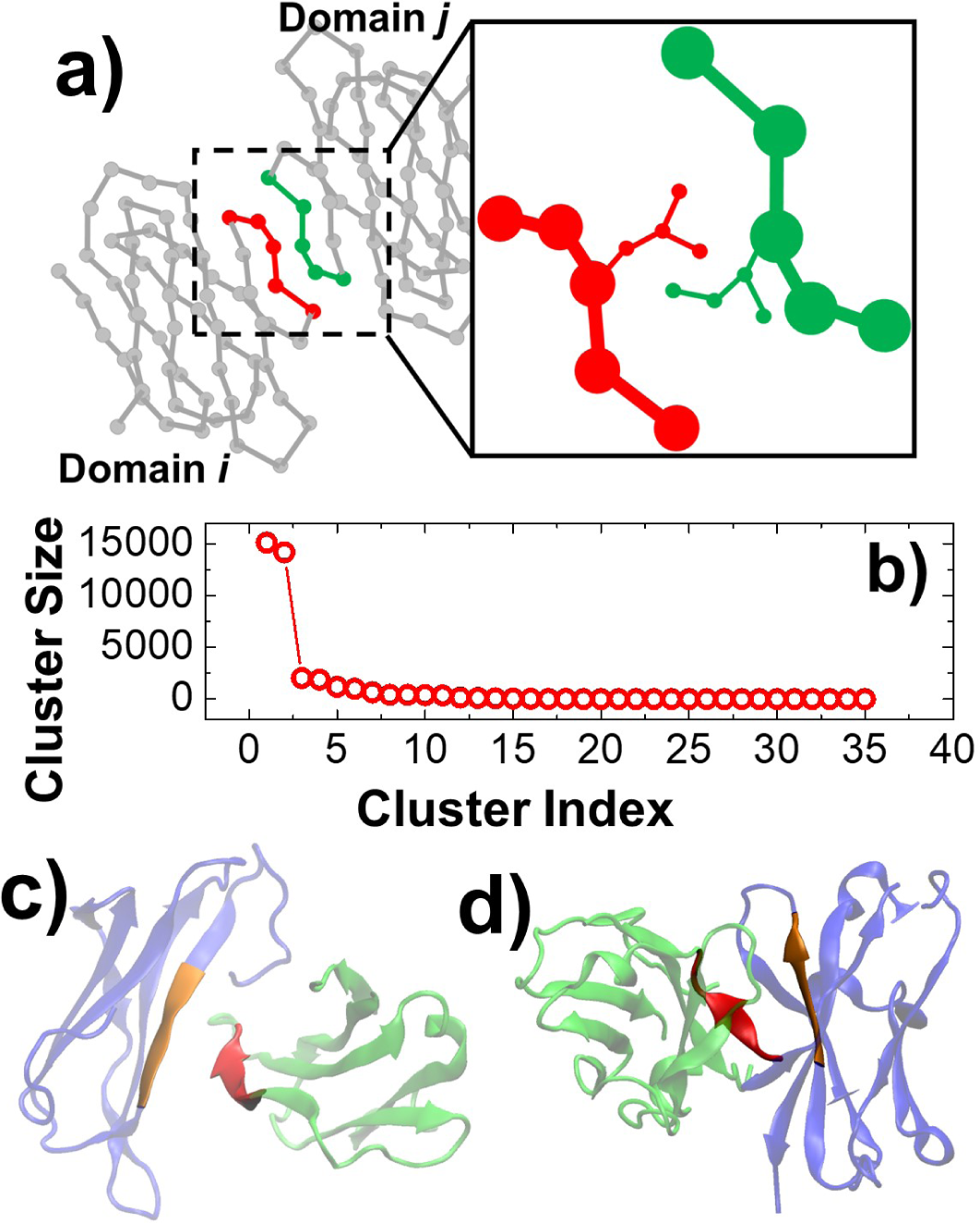
We defined interface fragment pairs from two interacting Ig domains **(a)**. One fragment is from domain *i* and the other one is from domain *j*. Additionally, the sidechain of the residue in the middle of one fragment form at least one atomic contact with the sidechain of the residue in the middle of the other fragment. We generated 37968 interface fragment pairs from 831 Ig domain interactions according to above definition, and clustered them into 36 groups based on their structural similarity. The numbers of interface fragment pairs in all groups were ranked, as shown in **(b)**. The structures of these fragment pairs in the two dominant groups are plotted in **(c)** and **(d)**, respectively. Two interface fragments are colored in red and orange, while their corresponding domains are colored in green and blue.

We classified all derived interface fragment pairs based on their structural similarity. The root-mean-square-difference (RMSD) of the Cα atoms between two comparing fragment pairs is used as the criterion of structural similarity. Detailed clustering algorithm is introduced in the **Materials and Methods**. Using 3 Angstrom as the RMSD cutoff value in the clustering, we found that all 37968 fragment pairs can be classified into only 36 different structural groups. We further counted the number of fragment pairs in all groups and ranked them in decreasing order. The results are shown in **Figure 2b**. Interestingly, the figure suggests that conformations of interface fragment pairs are dominated by the first two clusters. These two clusters together contain more than 75% interface fragment pairs in the library. The representative structures of interface fragment pairs from these two most abundant clusters are thereby plotted in **Figure 2c** and **2d**, respectively. Two fragments in the plots are colored in red and orange, while the interacting Ig domains that these fragments belong to are colored in green and blue. The fragment pairs in the first cluster correspond to two interacting β-strands that are antiparallel to each other (**Figure 2c**). On the other hand, the fragment pairs in the second cluster correspond to two interacting β-strands that are parallel to each other (**Figure 2d**).

Taken together, a library of 37968 interface fragment pairs were constructed from a non-redundant structure database of Ig domain interactions. The fragment pairs in the library were used as starting point to predict interactions between two cell surface proteins which contain extracellular Ig domains, as described in the next section.

### 2.2. Develop a machine learning framework to identify protein-protein interactions based on the interface fragment pair library

In order to identify potential interactions between two cell surface proteins which contain extracellular Ig domains, the information embedded in the interface fragment pair library will be fed into a machine-learning-based predictor. The procedure of developing this machine learning framework is as follows. We first generated sequence profiles for all 37968 interface fragment pairs in the library. The motivation of constructing the sequence profiles is to link all proteins that are evolutionarily related to the Ig domains in our database. In order to do so, PSI-BLAST [44] was carried out to all primary sequences of Ig domains in the non-redundant structure database. This is done with three iterations of searching against non-redundant (NR) sequence database. An E-value threshold of 0.0001 was used, which is a typical value adopted in previous studies [45]. After PSI-BLAST, position-specific scoring matrix (PSSM) for all residues on the corresponding interface fragment pairs were saved. As shown in **Figure 3a**, the derived sequence profiles for a specific interface fragment pair are two matrices with 5 columns and 20 rows. In these 5×20 matrices, the number of columns corresponds to the length of each fragment that contains 5 residues. The number of rows in the matrices represents all 20 types of amino acids. Each unit in the matrixes is a score collected from the PSSM matrix, indicating how likely if the original residue in the fragment will be replaced by the relative other type of amino acid.

**Figure 3:**
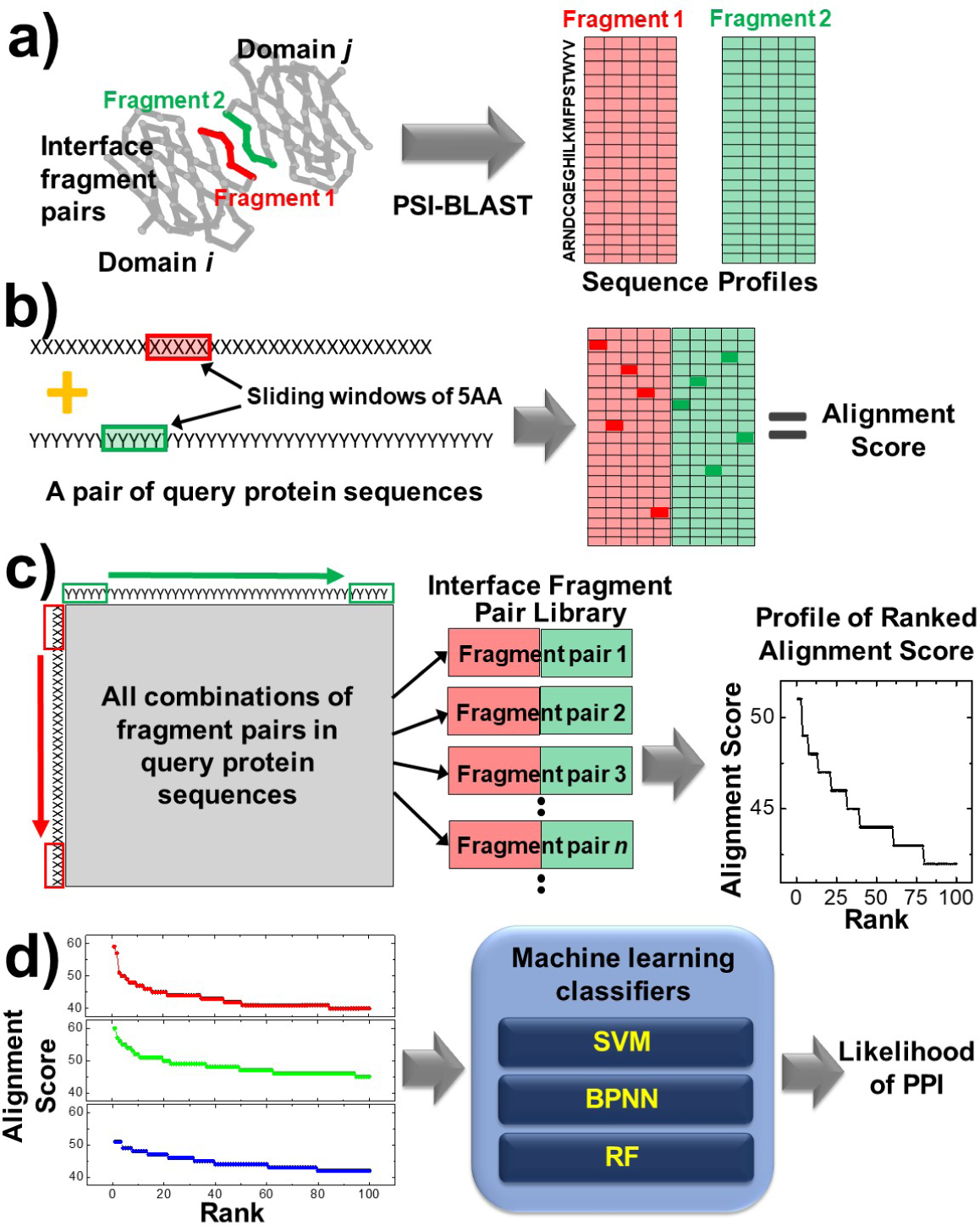
The 20-dimension sequence profiles for a specific interface fragment pair were generated from three iterations of PSI-BLAST, as shown in **(a)**. We then assigned a sliding window to each sequence in a pair of query proteins. We can align windows in both proteins to the sequence profiles of any interface fragment pair in the library and an alignment score can be calculated, as shown in **(b)**. By moving both windows from N-terminal to C-terminal and calculating alignment scores for each of these combinations to all interface fragment pairs in the library, a profile of ranked alignment score can be generated for a specific pair of query proteins. One example of the alignment score as the function of rank is shown in **(c)**. As the final step shown in **(d)**, when profiles of different pairs of query proteins are fed into multiple machine learning classifiers, after training, the likelihood of whether these proteins can form interactions will be derived.

For a pair of query proteins, we want to know if they can potentially form an interaction based on their primary sequences. We assigned a sliding window to each query protein. The length of the sliding window equals 5 amino acids, the same as the length of interface fragment pairs. When the sliding windows are located at specific positions of the query proteins, we can align them to the sequence profiles of any interface fragment pair in the library, as shown by the schematic in **Figure 3b**. An alignment score will be then calculated as follows. There are two different scenarios to align the two sliding windows to the sequence profiles of an interface fragment pair. As the first scenario, the red window can be aligned to the red matrix and the green window can be aligned to the green matrix. Otherwise, the red window can also be aligned to the green matrix and the green window can be aligned to the red matrix as the second scenario. For either scenario, we will map each of the five residues in one sliding window to the sequence profiles of the corresponding fragment. For instance, if the first residue in the window is Aniline, we will look up the first column in the 5×20 matrix and find out the value of the matrix unit which corresponds to the Aniline. Similarly, if the third residue in the window is Lysine, we will look up the third column and find out the value which will be cost to replace the original residue in the interface fragment to Lysine. The score will be added up from the first to the last residue in the window. Following the same procedure, the sliding window in the second query protein will be aligned to the sequence profiles of the other interface fragment and the score will the calculated. Both alignments will be added into a total score. The total score calculated from the first scenario is different from the total score calculated from the second scenario. Because the alignments in both scenarios are independent of each other, the final alignment score will be determined by adopting the higher total score from these two scenarios.

Given the calculation of alignment score for sliding windows, all combinations of fragment pairs in a given pair of query proteins were evaluated by moving both windows from N-terminal to C-terminal of their corresponding sequences, as illustrated in **Figure 3c**. For each specific combination, the residues in two sliding windows were aligned to the sequence profiles of all 37968 interface fragment pairs. The scores of all alignments were calculated. The highest alignment score was then saved for that specific combination. We ranked the highest alignment scores for all possible combinations between two query protein sequences. The top 100 highest alignment scores were selected. A profile of ranked alignment scores (PRAS) was finally generated by ranking these top highest alignment scores into decreasing order. One example of PRAS is represented by the plot on the right side of **Figure 3c**. The pattern of this profile was used as the fingerprint for a specific pair of query protein sequences. Various machine learning algorithms were implemented to recognize these patterns, as described below.

The left column of **Figure 3d** shows examples of PRAS for three pairs of cell surface proteins. The plot with red curve shows that the top 2 alignment scores are significantly higher than the rest in the profile. The plot with green curve shows the overall level of all alignment scores is high, while the plot with blue curve shows the opposite where the overall level of all alignment scores is low. All these pieces of information were then sent to the machine learning classifiers. We expect that the machine-learning-based models can capture the patterns hidden in these different profiles, and further detect the signals which indicates interactions between proteins. In detail, three different classification algorithms of machine learning were applied. They are: 1) support vector machine (SVM); back-propagation neural network (BPNN); and random forest (RF). The inputs of these classifiers are the profile of ranked alignment scores constructed for a specific pair of query protein sequences. It is a vector with 100 dimensions. The binary output from each classifier indicates whether two query protein can interact with each other or not. A consensus voting strategy was further employed to make an integrative decision from the outputs of three classifiers. More detailed description of these classification methods can be found in the **Materials and Methods**. In the next section, we trained these machine learning classifiers against an experimentally measured extracellular interactome of human Ig cell surface proteins.

### 2.3. Benchmark the machine-learning-based method against an experimentally-derived human interactome of cell surface proteins

A high-throughput experimental platform has recently been developed. The platform is based on the combination between an ELISA-based extracellular binding assay and an automated pooled-protein strategy [33]. The method was applied to screen the interactions among human secreted and single transmembrane cell-surface proteins in Ig superfamily. A total number of 426 PPIs between 564 proteins were detected, most of which have not been reported before but been confirmed later by other experimental methods. We compared these interactions to the ones generated from another PPIs prediction method, PrePPI [46]. PrePPI is currently one of the most advanced computational platforms that predicts protein-protein interactions. A Bayesian framework that integrates different sources of experimentally measured or computationally modeled information, including protein structures, functions, and evolutions, was designed to determine the likelihood of interactions in the human proteome. It has been used to predict Ig and Cadherin domain interactions [47]. A total of 8919 interactions were predicted by PrePPI among the same 564 cell surface proteins in human. However, only 77 of them overlap with the interactions detected by above high-throughput experimental platform. Therefore, a method that specifically focuses on the extracellular interactions between cell surface proteins is necessary. Here, we used the PPIs information derived from the experimental platform as a benchmark set to train our machine learning classifiers. Two different strategies were adopted to test the performance of our benchmarking results, as described below.

The first strategy consists of 10 runs of leave-one-out cross-validations, as shown by **Figure 4a**. Within each run, we randomly selected 500 pairs of proteins from the benchmark set. Among these 500 pairwise combinations, 250 of them are positive data. These 250 pairs of proteins were randomly selected from the 426 PPIs which interactions have been successfully identified in the above experimental study. The remaining 250 pairs of proteins were negative controls. They were randomly selected from the pairs of proteins whose interactions were not detected in the above experiment using the combination of an ELISA-based extracellular binding assay and an automated pooled-protein strategy [33]. We built the PRAS profiles for all 500 pairs of proteins based on the sequences in their extracellular regions. If both sequences of an interface fragment pair in the library share higher than 90% of sequence identity with these 500 pairs of proteins, it was removed from the profile construction. The profiles of both positive and negative data were then sent to the machine learning classifiers. A total number of 500 iterations were carried out for each run of leave-one-out cross-validations. Within each iteration, one of the 500 protein pairs was selected for test, while the remaining 499 pairs were used for training. The 100-dimensional profiles of the 499 protein pairs were fed into the classifiers as the inputs for training, while their expected outcomes about whether a corresponding protein pair can form interaction or not were used as the targeted outputs for training. Afterwards, the profile of the one testing protein pair was fed into the trained classifiers, and prediction of interaction was made. Predicted results for all protein pairs were collected after these 500 leave-one-out iterations.

**Figure 4:**
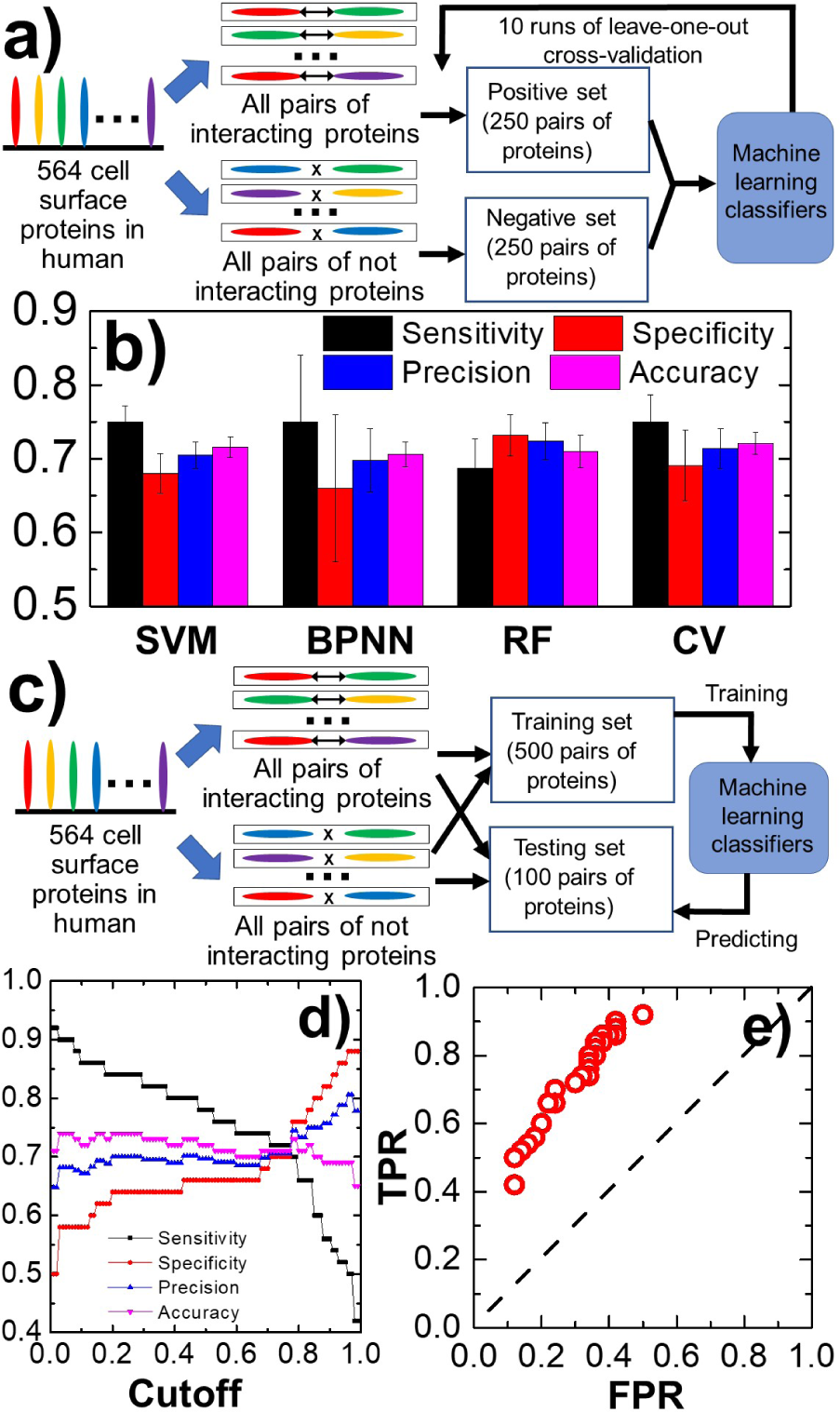
We tested our machine learning methods to an experimentally derived dataset which contains 564 cell surface proteins in human using two different strategies of cross-validation. In the first strategy **(a)**, 10 runs of leave-one-out cross-validations were carried out. The average sensitivity, specificity, precision and accuracy were calculated to calibrate the performance of cross-validations. The cross-validation results are summarized in **(b)** as histograms. In the second strategy **(c)**, an independent testing set was randomly selected in addition to the training set. After 10 runs of training, the probability of interaction was calculated for each protein pair in the testing set. Given the predicted probability, a threshold of interaction was defined above which a given pair of proteins can be predicted as positive. The calculated sensitivity, specificity, precision and accuracy were plotted in **(d)** as we increased the threshold from 0 to 1. We also plotted the correlation between true positive rate and false positive rate based on the prediction results of different threshold values, as shown in **(e)**.

The performance of the prediction was evaluated by calculating the sensitivity, specificity, precision and accuracy from the collected data which are defined in the **Materials and Methods**. After 10 runs of cross-validations, the calculated results were further averaged. The overall performance of different machine learning models is summarized in **Figure 4b**. Specifically, the black, red, blue and purple bars in the histograms represent the averaged values of calculated sensitivity, specificity, precision and accuracy, respectively. Different types of classifiers are indicated in the bottom. The figure shows that different models can attain similar results. The overall sensitivity, specificity, precision and accuracy in these classifiers are all around 70%. Additionally, the consensus voting based on the outputs from the other three machine learning algorithm can archive the highest accuracy of 72%.

In the alternative strategy (**Figure 4c**), 100 pairs of proteins were randomly selected as an independent testing set. Among these selected protein pairs, 50 of them have validated interactions and were treated as positive data, while the rest 50 pairs were chosen as negative control. In addition to the testing set, a training set consisting of 500 protein pairs was constructed, in which half of them are positive for interactions and the other half are negative. Moreover, none of the protein pairs in the training set is included in the testing set. We built the PRAS profiles for all pairs of proteins in the training and testing sets. If both sequences of an interface fragment pair in the library share higher than 90% of sequence identity with any protein pair in the training or testing sets, it was removed from the profile construction. The profiles for all protein pairs in the training set were first sent to the machine learning models. After training, the profiles of testing set were input to the model to generate the predictions. This process was repeated for 10 times. Within each run, a different training set with 500 randomly selected protein pairs was constructed but the testing set remained unchanged. After all 10 runs of cross-validations, the probability of interaction was calculated for each of the 100 protein pairs in the testing set as ∑δ(i,j)/N, in which δ(i,j) is the output of classifier i from the j^th^ run and the value of output is either 1 for positive or 0 for negative. There is a total of 4 different classifications and 10 independent runs of training. As a result, the value of N equals to 4×10=40.

Given the predicted probability for all 100 protein pairs, a threshold of interaction was defined. A pair of proteins will be predicted as positive to form an interaction if its calculated probability is above this threshold. Otherwise, it will be predicted as not forming an interaction if the probability is below the threshold. **Figure 4d** plots our prediction results as we increased the value of threshold from 0 to 1. The figure shows that along with the raise of threshold, the specificity (red) and precision (blue) increase from a low level, while the sensitivity (black) keeps decreasing. When the threshold is within a small window between 0.7 and 0.8, the optimal performance of the system was obtained in which all specificity, precision, sensitivity and accuracy are above 70%. We also analyzed the correlation between true positive rate (TPR) and false positive rate (FPR) from the prediction results. TPR and FPR are defined in the Materials and Methods. These two rates are collectively changed under different values of threshold, which form a receiver operating characteristic (ROC) curve [48] as shown by the red circles in **Figure 4e**. The area under the curve (AUC) score calculated from the ROC curve equals 0.76. In comparison, the dashed diagonal line, also known as the line of no-discrimination, indicates that the prediction is completely generated from random guess. The large space between ROC curve and the line of no-discrimination suggests our benchmark tests are significantly better than random on a statistical level.

We compared our results with a simple classifier in which only the highest alignment score in the PRAS profile was used for prediction. We randomly selected 100 pairs of proteins to test the model. Among these selections, only 50 of them have experimentally validated interactions. During the test, the interaction between two proteins was predicted to be positive, if their highest alignment score was above a cutoff value. On the other hand, if the highest alignment score was below the cutoff, the interaction was predicted to be negative. The cutoff was selected so that the number of positive predictions equaled the number of negative predictions. We repeated this analysis five times and calculated the accuracy of the prediction within each time. The average prediction accuracy of this simple classifier equals 61±2%. This accuracy is better than random guessing which is around 50%, but still significantly lower than the results from our machine-learning-based models. This test indicates that the highest alignment score contains a certain degree of information about the relation of two selected proteins, but not enough to capture the full signals of their interaction. Therefore, it is necessary to develop the higher-dimensional PRAS profiles and apply machine learning algorithms to recognize the patterns in these profiles.

In summary, the benchmark tests using both cross-validation strategies archived similar performance. The testing results suggest that our machine learning classifiers, after trained by the signals embedded in the interface fragment pair library, are pretty reliable to identify new potential interactions between cell surface proteins with extracellular Ig domains.

### 2.4. Apply the calibrated machine-learning classifier to predict the interactions of cell surface proteins in C. elegans on a systematic level

After the machine learning classifiers have been benchmarked by the interactome of human cell surface proteins, we applied the calibrated models to predict interactions between cell surface proteins in other species. Specifically, we selected 46 secreted or single transmembrane cell-surface proteins in Caenorhabditis elegans. All these proteins contain one or multiple extracellular domains that belong to either Ig superfamily or the superfamily of Ig-like fold. A full list of these proteins, including their gene names and number of Ig domains, can be found in the **Supporting Table 1**. We carried out predictions to all possible pairwise interactions among these proteins. A total number of combination equals 46×(46+1)/2=1081. We calculated the PRAS profiles for all the 1081 protein pairs based on the sequences in their extracellular regions as the inputs to machine learning classification. Ten runs of predictions were performed for each combination. Within each run, 500 pairs of human proteins were picked as training set. Half of the protein pairs in the training set were identified as positive for interactions and the other half are negative. After training, the PRAS profile of each protein combination was fed into all four machine learning models for prediction. When all 10 runs of prediction completed, the probability of interaction can be derived for a specific protein pair based on the equation ∑δ(i,j)/N as introduced in the last section.

A threshold of 0.7, which gave the most optimal performance during the benchmark test, was used to determine whether two proteins can form an interaction or not. If the calculated probability of interaction for a protein pair is higher than this threshold, we claim that these two proteins can interact with each other. On the other hand, the calculated probability of interaction for another protein pair which is lower than this threshold indicates that these two proteins cannot interact with each other. Following this criterion, a total number of 204 positive interactions have been predicted among all 1081 combinations. A network representation of all predicted interactions is illustrated in **Figure 5a**. The nodes in the plot denote all proteins involved in these interactions with their gene names marked on the corresponding nodes, and the edges represent all predicted interactions. We found 32 out of 46 proteins form at least one interaction with others, while binding partners of the remaining 14 proteins cannot be detected by our machine learning predictions. In order to further characterize the topology of our constructed network, we counted the numbers of binding partners for all proteins. They were plotted as a histogram in **Figure 5b**. The figure shows that more than one third of proteins have less than 10 binding partners. In contrast, there is one protein, zig-6, can interact with all proteins in the network including itself. Zig-6 is a secreted protein containing two Ig domains. It is expressed in body wall musculature; enteric muscle; head neurons; and vulval muscle, and was found to be involved in homeostatic process [49]. Unfortunately, its interactions with other protein in C. Elegans have not been previously documented.

**Figure 5:**
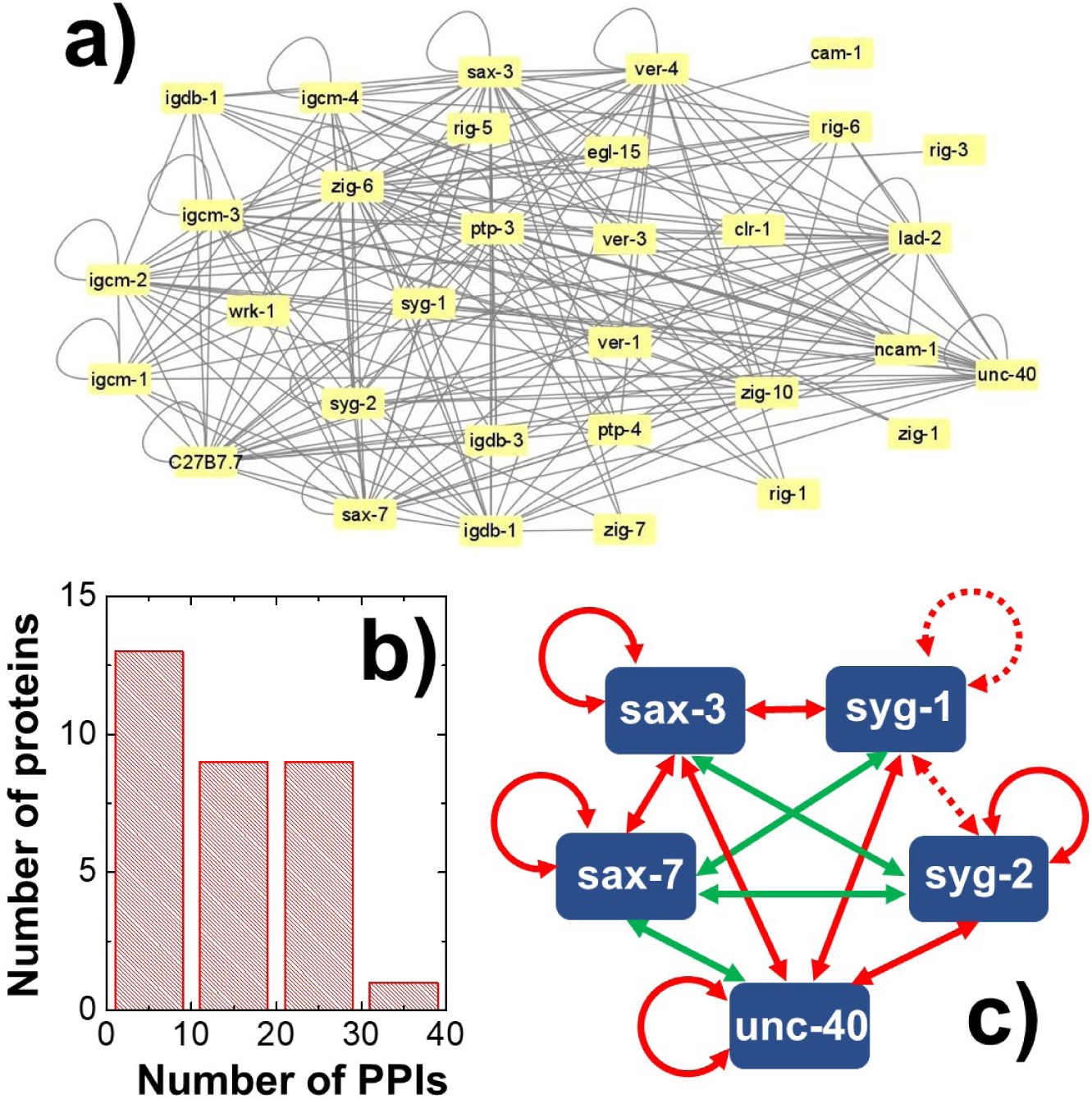
After we trained our machine learning classifiers against the interactome of human cell surface proteins, we applied it to predict interactions between cell surface proteins in C elegans. A total number of 46 secreted or single trans-membrane proteins from C elegans were selected, which contain one or multiple extracellular domains that belong to either Ig superfamily or the superfamily of Ig-like fold. We screened every possible interaction between these proteins. As a result, a total number of 204 positive interactions have been predicted. A network representation of these interactions is illustrated in **(a)**. As shown in **(b)**, the number of binding partners for each protein in the network was counted. Finally, a subset of 5 proteins were picked. These proteins are relatively well studies and their interactions have been well documented. A network diagram of interactions among these 5 proteins is illustrated in **(c)**, in which 9 out of 13 predicted interactions (red solid arrows) have been discovered in previous experimental studies. The other 4 interactions are firstly reported in this study, as represented by green solid arrows. Another two interactions (red dashed arrows) have been previously observed but cannot be predicted by us.

In order to validate our predictions, we selected a subset of proteins that are relatively well studied. Specifically, the subset consists of 5 proteins. They are: syg-1, syg-2, sax-3, sax-7 and unc-40. Syg-1 and syg-2 are cell adhesion molecules necessary for synaptic specificity [50]. Unc-40 is enriched at presynaptic sites and was found to play essential roles in regulating synaptogenesis [51]. Receptors sax-3 and sax-7 are involved in the stimulation of axon outgrowth and maintenance of neuronal positions [52]. Because the functions of these proteins are highly overlapped, it is reasonable to assume that they can form various homophilic and heterophilic interactions among each other. As a result, a network diagram of our predicted interactions among these 5 proteins is illustrated in **Figure 5c**. In details, our machine learning models predicted a total number of 13 interactions in the selected subset. Among these 13 interactions, 9 of them have been documented in previous experimental studies. These 9 interactions are marked by red solid arrows in the network. Detailed references of these experimental studies can be found in the **Supporting Table 2**. The rest 4 interactions, as marked by the green solid arrows in the network, are newly identified by our predictions. These interactions proposed by our computational study can be experimentally verified by others in the future. Finally, two other interactions, the homophilic interaction between syg-1 and the heterophilic interaction between syg-1 and syg-2, have been previously observed but cannot be predicted by us, as marked by the red dashed arrows in the network. It is worth mentioning that the signal of interaction between syg-1 and syg-2 has been partially captured by our prediction. However, the predicted binding probability equals 0.625, right below the threshold.

Our study also shows that there are 14 protein which have no interaction with any others. One example is oig-1, which is a single-Ig-domain secreted protein [53]. It is known to be involved in regulating synapse formation in C. elegans. The protein is co-expressed in connected neurons along with at least three other proteins in our test set: rig-6, sax-3, and sax-7. It is hard to imagine how this protein can carry out its function without interacting with others. It is thereby interesting to understand why there was not positive signals generated from our classifiers for this protein. We speculate this is possibly due to the following two reasons. Firstly, the signals our method used to detect potential interactions is originated from the information of homologous sequence in the interface fragment pair library. It is likely that oig-1 and its Ig-domain binding partners interact with each other through the interfaces from which the structures of relevant homologous proteins have not been experimentally derived. The potential of how to improve our predictions by using more remote information in interacting proteins is discussed in the next section. It is also possible that oig-1 carries out its functions by interacting with proteins that only contain domains in other superfamilies. Its interactions with other Ig proteins therefore cannot be detected. It is worth mentioning that the interactions between proteins in Ig and other superfamilies have been previously observed. For instance, Liu et al. found that Ig proteins BTLA/CD160 can bind to HVEM, a member in the tumor necrosis factor receptor (TNFR) superfamily [54].

## Concluding Discussions

Many cell surface proteins are promising therapeutic targets due to their essential functions in regulating intercellular communications within many physiological or pathological processes. However, most current high-throughput screening techniques are less sensitive to detect the extracellular interactions between these proteins. Although the advancements in method development of machine learning enable us to computationally predict protein-protein interactions with a more convenient way, its application to cell surface proteins has not been documented before. In order to reach this goal, we developed a new computational framework by specifically focusing on the interactions between proteins with domains in Ig fold. A non-redundant structural database of interactions between Ig domains was constructed and transformed into a fragment pair library. For a given pair of query protein sequences, a high dimensional profile was generated based on the information collected in the library. This profile was then sent to various machine-learning-based predictors which was trained by an experimentally interactome of cell surface proteins in human. Cross-validation results show that our method can achieve higher than 70% accuracy. We then applied this method to systematically predict the interactions of 46 cell surface proteins in C elegans. For a small subset of well-studied proteins, most of the interactions predicted by our method have been experimentally confirmed in the literature. Taken together, our computational platform serves a useful tool to help identifying potential new interactions between cell surface proteins in addition to current state-of-the-art experimental techniques.

As we showed in the results, for some proteins of interest, our current method was not able to detect any positive signal for their potential binding partners. We applied PSI-BLAST to generate sequence profiles for all interface fragment pairs in the library and used them to construct the feature space in machine learning classifications. As a result, it might be difficult for query proteins which share little information of sequence homology with Ig domains in the library to be captured by the classifiers after training. This issue can be improved in the future by integrating profiles generated not only by PSI-BLAST, but also by structure comparison. These structure-based profiles can be constructed by adding information of secondary structure types and solvent accessible surface [55], or by comparing the conformations of local fragments from the proteins in our current database to those similar fragments in the entire PDB. The second alternative way to improve our method is to implement more advanced deep learning classification algorithms. The combination of sequence and structure profiles will lead to a higher dimensional feature space. Predictions based this new feature space could be benefitted by the application of deep learning methods such as convolution neural network (CNN) [56]. However, either the integration of new features from structural comparison or the application of deep learning methods require a higher number of data points during training to avoid model overfitting. One possible solution is to construct a larger benchmark dataset. For instance, an experimentally measured interactome of Drosophila cell surface proteins [57] could be added to the original dataset of human cell surface proteins in the future. Finally, there is a well-established interaction in C. elegans cannot be predicted by our method: the interaction between rig-5 and zig-8. We noticed that the crystal structure of heterodimer between N-terminal Ig domains of rig-5 and zig-8 was very recently deposited to PDB [58]. As a result, the signal of interaction between rig-5 and zig-8 has been missing in our database. This provides another avenue to improve our prediction by updating our database with the newest structural information of Ig domain interactions.

In reality, a large portion of cell surface proteins in our study contain multiple Ig domains in their extracellular regions. However, not all these domains are directly involved in protein-protein interactions [59]. For instance, type-I classic cadherins contain 5 extracellular domains, but only the N-terminal domains are responsible for their *trans*-dimerization. In order to understand the functions of cell surface proteins, it is not only necessary to identify whether they form interactions, but also more important to determine which specific domains in these proteins are engaged in binding [60]. The primary target of current method developed in this study is to predict if two proteins can form an interaction or not. However, it can be extended to predict the interactions between two specific Ig domains. In practice, instead of using the entire sequences of two query protein as inputs, we will narrow down the inputs to the sequences of individual Ig domains from two cell surface proteins, based on which we can use machine learning to predict whether these two domains bind to each other or not. Moreover, the non-redundant structural database of Ig domains interactions itself can be used as the benchmark set to train the machine learning classifiers. Using the above-mentioned classic cadherins as an example, all 5×5 pairwise combinations can be tested between their five extracellular domains. As a result, given the fact that two cadherins form a homophilic interactions, we will further be able to identify which domains are responsible for their binding. The expected prediction output is that the binding probability between two N-terminal domains would be much higher than other combinations. If success, our prediction will generate a shortlist of domain-domain pairs between two interacting cell surface proteins for experimental verification. This can greatly reduce the complexity of experimental tests in which all possible domain combinations will have to be enumerated by brute-force screening.

Future extension of current study also includes applying our computational models to predict the interactions between domains in other superfamilies. One example is the domains in death domain (DD) superfamily [61]. Many protein complexes involved in inflammatory signaling pathways are assembled through the homotypic interactions between domains that belong to this superfamily. It comprises of the DD subfamily [62, 63], the death effector domain (DED) subfamily, the caspase recruitment domain (CARD) subfamily and the pyrin domain (PYD) subfamily, which all share a similar fold of six-helical orthogonal bundle [64]. Interestingly, not all members in the DD superfamily are observed in complexes. A small portion of them exist as monomers, while other domains can only form homophilic or heterophilic interactions across different subfamilies. For instance, three types of DD domains, MyD88, IRAK2 and IRAK4 can be aggregated together into a helical oligomer, called Myddosome [65]. The application of our machine learning model to domains in the DD superfamily by constructing the sequence profiles of fragments at their binding interfaces will be helpful to predict whether a DD domain with unknown function can be assembled into complexes or not.

## 3. Materials and Methods

### 4.1 The construction of non-redundant structural database for Ig domain interactions

In order to build a database specifically for Ig-fold domains, we first collected information about domain-domain interactions from iPfam [66]. There is a list of more than 2×10^5^ records of domain-domain interactions in iPfam, including both intramolecular and intermolecular homophilic or heterophilic interactions. An interaction was called by iPfam when one or more bonds are formed between two protein domains. Bonds are defined by previous studies [67], which are based on both chemical and geometric properties of the atoms involved in the molecules. For each record, the Pfam id numbers of both interacting domains are presented. Because Ig-fold domains in almost all cell surface protein are found to be from two Pfam clans: the Ig superfamily (CL0011) and the Ig-like fold superfamily (CL0159). We searched the Pfam id in all records of domain-domain interactions in iPfam again these two clans. Specifically, domains in clan CL0011 consists of 28 Pfam protein families, while domains in clan CL0159 contains 82 Pfam protein families. The Pfam id of all these 110 protein families can be found in the **Supporting Table S3**. During our search, if both interacting domains in an iPfam record are from one of these 110 protein families, the relative information, including the primary sequence of each domain and the PDB id, will be saved for the next step. As a result, a total number of 3829 interactions were obtained between domains in Ig superfamily or Ig-like fold superfamily.

There is redundant information in the 3829 pairs of Ig domain interactions. For instance, two records of interactions might be from the same pair of proteins, but their structures were determined by different groups therefore have different PDB ids. We removed this redundancy from the database in order to avoid the statistical bias. In detail, we compared the sequences for all 3829 pairs of protein domains. Any two pairs of interactions in which both domains share higher than 90% of sequence identity are not allowed in the database. In another word, if the first domain in pair *i* shares higher than 90% of sequence identity with the first domain in pair *j*, while the second domain in pair *i* shares higher than 90% of sequence identity with the second domain in pair *j*, either one of these two pairs of interactions has to be excluded from the database. On the other hand, if the first domain in pair *i* shares higher than 90% of sequence identity with the first domain in pair *j*, but the second domain in pair *i* shares low sequence identity with the second domain in pair *j*, these two pairs can coexist in the database. After we applied the inspection of redundant information, 831 pairs of Ig domain interactions remain in the database. Finally, the atomic coordinates for all these 831 interacting domains were downloaded from PDB.

### 4.2 Structural based clustering of interface fragment pairs

As introduced in the **Results**, we defined interface fragment pairs for residues that are located at the binding interfaces of two interacting Ig domains. All these interface fragment pairs were collected by scanning the structure of each Ig domain interaction in the non-redundant database. The similarity between different interface fragment pairs was compared on a systematic level by a clustering procedure. Specifically, an initial interface fragment pair was randomly picked from the library as the first cluster. For the next randomly picked fragment pair, structural comparison with the first cluster was carried out by calculating the RMSD between two pairs. If the RMSD is smaller than a predetermined threshold, the latter pair will be included into the first cluster. Otherwise, the second cluster will be created. This procedure was iterated. Assuming that, before the entry of the *i^th^* picked fragment pair, the previous *i-1* pairs have already formed *m* clusters. Then the newly picked pair will be compared to all the previous *i-1* pairs and find its nearest neighbor with the lowest RMSD. If this lowest RMSD value is larger than the threshold, we will create the *m+1* cluster for this entry. Otherwise, it will be assigned to the same cluster its nearest neighbor belongs to.

In order to test the impact of the RMSD cutoff on the clustering results, we changed it to different values. For each cutoff value, clustering was carried out over all 37968 interface fragment pairs. **Figure S1** shows the histograms of the derived cluster numbers as a function of the RMSD cutoff from 1 Angstrom to 5 Angstrom. The figure indicates that the total number of clusters increases quickly when the cutoff value becomes lower. If the cutoff equals 5 Angstrom, all fragment pairs can be clustered into one group. As a result, clustering loses its sensitivity under this large cutoff value. On the other hand, when the cutoff equals 1 Angstrom, there are 16108 clusters. As a result, the interface fragment pairs within each cluster lose their structural diversity under this small cutoff value. After balancing diversity with sensitivity, a RMSD threshold of 3 Angstrom was adopted, which leads to a total number of 36 clusters.

### 4.3 Description of various machine learning algorithms used in the study

Our machine learning classifications are based on three algorithms. The feedforward back-propagation neural network (BPNN) was first implemented [68]. The input neurons of the network consist of 100 dimensions, the same as the PRAS profile. The output of the neural network is in the binary format. The network further contains a single hidden layer with four neurons. A sigmoid activation function was adopted. Weight of each neuron is modified using the back-propagation learning algorithm with a sum of square error function. The magnitude of the error sum in the learning process is monitored in each cycle. The learning is terminated when the network converges. Secondly, an iterative algorithm of the support vector machine (SVM) classifier was implemented [69]. The algorithm converts the classification into a problem of computing the nearest point between two convex polytopes. The Gaussian kernel was adopted in the model. The optimal values of all tunable hyper-parameters were determined by the grid search method. Finally, the random forest (RF) classifier was implemented based on the growth of many single decision trees [70]. Each single decision tree was built by using a number of randomly selected features. The total number of features is the dimensions of our input vectors corresponding to the profiles of top 100 ranked alignment scores. After the tree construction, a new data point can be traversed through each tree in the forest from the root to one of its leaf nodes, from which the class of the data point can be determined. Consequently, the algorithm chooses the classification that has the most votes over all the trees in the forest. Specifically, we allowed 500 trees to be grown in the forest. Two randomly selected features were used to split on at each node of trees. Values of parameters for all machine learning algorithms were selected to ensure that the training processes were stable.

A consensus voting (CV) strategy was further proposed to make an integrative decision from the outputs of above three machine learning classifiers [71]. Specifically, the integration of machine learning outputs for a pair of query proteins was derived by calculating *∑wi×δi*, in which *i* is summated through all three classifiers. The delta function *δi* is the binary signal generated from the *i^th^* machine learning classifier, which equals 1 for positive prediction, and -1 for negative prediction. The parameter *w_i_* introduces the weight of each machine learning algorithm during voting, and thus represents the relative contribution of each classification to the final consensus prediction. The positive or negative output from the voting corresponds to whether two input query proteins can form an interaction or not.

### 4.4 Evaluation of cross-validation performance

During cross-validation, each machine learning classifier will generate a binary signal for each input pair of protein sequences. Afterwards, the total numbers of true positive (TP), false positive (FP), true negative (TN), and false negative (FN) from these binary outputs were counted in order to calibrate the performance of our cross-validation results. A TP is marked for a pair of proteins which form an experimentally detected interaction, if our prediction also indicated that these two proteins can interact with each other. On the other hand, a FP is marked for a pair of proteins which interaction cannot be experimentally detected, but our prediction indicated that these two proteins can interact with each other. Similarly, a TN is marked for a pair of proteins which interaction cannot be experimentally detected, while our prediction also indicated that these two proteins cannot interact with each other. Finally, a FN is marked for a pair of proteins which form an experimentally detected interaction, but our prediction indicated that these two proteins cannot interact with each other.

Given these definitions, the overall sensitivity of cross-validation can be calculated by TP/(TP+FN). The specificity and precision were estimated by calculating the values of TN/(TN+FP) and TP/(TP+FP), respectively. Finally, the overall accuracy of the cross-validation results was derived by calculating the value of (TP+TN)/(TP+TN+FP+FN). As shown in the **Results**, we also calculated the true positive rate and false positive rate in order to plot the ROC curve. By definition, the true positive rate is equivalent to sensitivity. The false positive rate was derived from the ratio of FP to the combination of FP and TN

## Acknowledgement

This work was supported by the National Institutes of Health under Grant Numbers R01GM120238 and R01GM122804. The work is also partially supported by a start-up grant from Albert Einstein College of Medicine. Computational support was provided by Albert Einstein College of Medicine High Performance Computing Center.

## Author Contributions

S.E. and Y.W. initiated the research; Z.S and Y.W. designed the research; Z.S. performed the research; Z.S., B.G., S.E. and Y.W. analyzed the data; Z.S. drafted the paper; S.E. and Y.W. revised the paper.

## Data Availability

All data generated from this study and the relevant source codes of the machine learning models can be found in the **Supplemental Document** or in the GitHub repository https://github.com/wulab-github/IgPPI.

