## Supplementary material for "Prediction of Interactions between Cell Surface Proteins by Machine Learning": IgPPIpred_SupportingInformatiojn_052023.pdf

### Supporting Information

Zhaoqian Su[1], Brian Griffin[2], Scott Emmons[2], Yinghao Wu[1,*]

[1]Department of Systems and Computational Biology, Albert Einstein College of Medicine, 1300 Morris Park Avenue, Bronx, NY, 10461

[2]Department of Genetics, Albert Einstein College of Medicine, 1300 Morris Park Avenue, Bronx, NY, 10461

| Gene Name | Domain Index |
|---|---|
| cam-1 (ROR) | Ig + Frz + Kr + TyrKinase |
| C27B7.7 | 3 Ig + 6 Fn3 (secreted) |
| clr-1 | 1 Ig + 2 Fn3 + TyrPhosphatase |
| egl-15(FGFR) | 3 Ig + TM + TyrKinase |
| igcm-1 | 7 Ig + 2 Fn3 + TM |
| igcm-2 | 3 Ig + 2 Fn3 + TM |
| igcm-3 | 3 Ig + TM |
| igcm-4 | 3 Ig + TM |
| igdb-1 | 1 Ig + 4 Fn3 + 2 DB + TM |
| igdb-2 | 2 Ig + 5 Fn3 + 4 DB + TM |
| igdb-3 | 1 Ig + 1 Fn3 + 2 DB |
| lad-2 (L1) | 6 Ig + 5 Fn3 + TM |
| ncam-1 | 5 Ig + 1 Fn3 + TM |
| oig-1 | 1 Ig (secreted) |
| oig-2 | 1 Ig (secreted) |
| oig-3 | 1 Ig (secreted) |
| oig-4 | 1 Ig (secreted) |
| oig-5 | 1 Ig (secreted) |
| oig-6 | 1 Ig + TM |
| oig-7 | 1 Ig + TM |
| oig-8 | 1 Ig + TM |
| ptp-3 (LAR) | 3 Ig + 9 Fn3 + TyrPhosphatase |
| ptp-4 | 1 Ig + 3 Fn3 + TyrPhosphatase |
| rig-1 | 6 Ig + 2 Fn3 + TM |
| rig-3 | 4 Ig + GPI |
| rig-5 | 3 Ig + GPI |
| rig-6 | 6 Ig + 4 Fn3 + GPI |
| sax-3 (Robo) | 6 Ig + 3 Fn3 + TM |
| sax-7 (L1) | 6 Ig + 5 Fn3 + TM |
| syg-1 | 5 Ig + TM |
| syg-2 | 8 Ig + 1 Fn3 + TM |
| unc-40(DCC) | 4 Ig + 6 Fn3 + TM |
| ver-1 | 5 Ig + TM + TyrKinase |
| ver-3 | 4 Ig + TM + TyrKinase |
| ver-4 | 4 Ig + TM + TyrKinase |
| wrk-1 | 3 Ig + 1 Fn3 + GPI |
| zig-1 | 2 Ig + TM |
| zig-10 | 2 Ig + TM |
| zig-2 | 2 Ig (secreted) |
| zig-3 | 2 Ig (secreted) |
| zig-4 | 2 Ig (secreted) |
| zig-5 | 2 Ig (secreted) |
| zig-6 | 2 Ig (secreted) |
| zig-7 | 2 Ig (secreted) |
| zig-8 | 2 Ig (secreted) |
| zig-9 | 2 Ig (secreted) |

**Table S1**: The list of cell surface proteins with Ig domains in C Elegans used as a test set in this study

| Gene 1 | Gene 2 | Method of determination | References |
|---|---|---|---|
| unc-40 | unc-40 | In vitro binding analysis | http://dx.doi.org/10.1038/nn956 |
| unc-40 | sax-3 | In vitro binding analysis | http://dx.doi.org/10.1038/nn956 |
| sax-3 | sax-3 | In vitro binding analysis | http://dx.doi.org/10.1038/nn956 |
| sax-3 | sax-7 | Co-IP, S2 cell aggregation | http://dx.doi.org/10.1016/j.devcel.2018.10.028 |
| sax-3 | syg-1 | Co-IP | http://dx.doi.org/10.7554/eLife.57921 |
| sax-7 | sax-7 | S2 cell aggregation | https://pubmed.ncbi.nlm.nih.gov/15775964/ |
| syg-1 | syg-1 | Co-IP | http://dx.doi.org/10.1371/journal.pone.0023598 |
| syg-1 | syg-2 | Co-IP, S2 cell aggregation, | http://dx.doi.org/10.1371/journal.pone.0023598 |
| syg-2 | syg-2 | Co-IP | https://doi.org/10.1371/journal.pone.0023598 |
| unc-40 | syg-1 | In vitro binding analysis | https://string-db.org/ |
| unc-40 | syg-2 | Gene co-expression | https://string-db.org/ |

**Table S2**: The experimental evidences of interactions between a subset of well-studied cell surface proteins in C Elegans

| Pfam Clan List | Pfam Family Index | | | |
|---|---|---|---|---|
| Ig superfamily (CL0011) | PF11465<br>PF09291<br>PF05326<br>PF16681<br>PF10430<br>PF16706<br>PF09085 | PF07679<br>PF13927<br>PF07686<br>PF13895<br>PF07654<br>PF00047<br>PF08205 | PF00207<br>PF03921<br>PF05790<br>PF16680<br>PF15910<br>PF08204<br>PF02440 | PF15028<br>PF02480<br>PF01537<br>PF02124<br>PF01688<br>PF16758<br>PF02960 |
| Ig-like fold superfamily (CL0159) | PF09067<br>PF07523<br>PF08770<br>PF13750<br>PF09134<br>PF16794<br>PF15886<br>PF03174<br>PF16407<br>PF16578<br>PF12690<br>PF10420<br>PF15418<br>PF11589<br>PF16820<br>PF08329<br>PF09238<br>PF07140<br>PF08191<br>PF09099<br>PF13752 | PF00041<br>PF00028<br>PF00630<br>PF01833<br>PF16184<br>PF14310<br>PF00801<br>PF02368<br>PF03168<br>PF01345<br>PF16656<br>PF08266<br>PF03443<br>PF02883<br>PF02494<br>PF05345<br>PF01835<br>PF07495<br>PF13205<br>PF08441<br>PF03404 | PF13573<br>PF13585<br>PF07705<br>PF03067<br>PF12245<br>PF16640<br>PF00927<br>PF13290<br>PF16655<br>PF16403<br>PF13860<br>PF16179<br>PF12733<br>PF04234<br>PF06280<br>PF01108<br>PF08752<br>PF02010<br>PF11614<br>PF09294<br>PF10029 | PF06312<br>PF07718<br>PF13629<br>PF11974<br>PF06832<br>PF13287<br>PF10633<br>PF07532<br>PF09240<br>PF08758<br>PF02369<br>PF02927<br>PF05753<br>PF01917<br>PF06030<br>PF06328<br>PF08428<br>PF16893<br>PF13753 |

**Table S3**: The list of Pfam families for domains in Ig fold used to construct our non-redundant structure database of Ig domain interactions

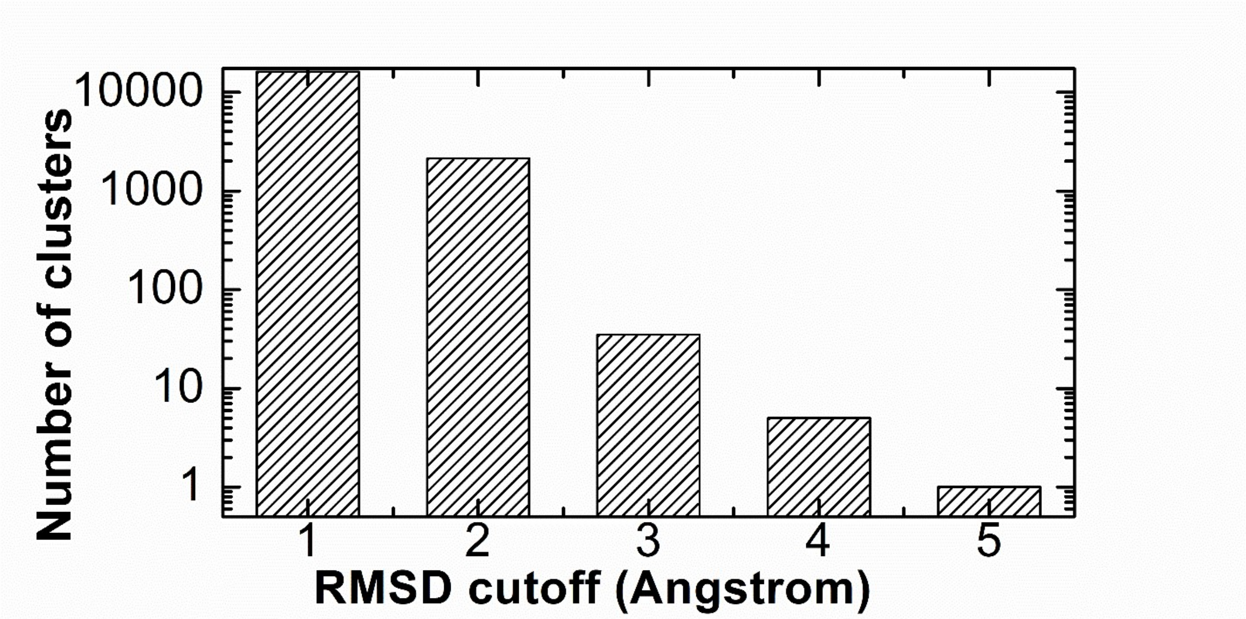


**Figure S1**: We clustered all the 37968 interface fragment pairs in our non-redundant structural database using different values of RMSD cutoff. The derived numbers of clusters under different RMSD values are plotted in the figure.
